# Maternal diesel exposure and maternal choline supplementation interactions in fetal and placental immune factors

**DOI:** 10.1101/2023.01.04.522511

**Authors:** Sara V Maurer, Jessica L Bolton, Staci D Bilbo, Christina L Williams

## Abstract

Air pollution causes widespread inflammatory changes in the body and brain. When exposure to air pollution occurs early in development, children exhibit impaired working memory ability (Sunyer et al., 2015). In addition, prenatal exposure to diesel particulate matter (DEP) increases inflammatory cytokine expression in the whole brain of embryonic day 18 (E18) males and leads to adverse long-term negative outcomes (Bolton et al., 2012). In contrast, dietary choline supplementation is negatively correlated with inflammatory cytokine production in adult rats and cultured human cells (Zhang et al., 2018; Jiang et al., 2014). When administered as a supplement to pregnant rats, choline also improves working memory in adulthood (Meck et al., 2008; Meck & Williams, 1999; 1997). The current study sought to determine if prenatal dietary choline supplementation protects against the effects of air pollution in the developing brain and in the placenta and fetal liver. These data revealed region-specific microglial morphology alterations in fetal brain and in inflammatory gene expression in the placenta and fetal liver (specifically, *Tnf, Tlr2, Tlr4*, and *Itgam*) due to maternal choline supplementation and/or maternal air pollution exposure. We found that DEP led to changes in microglial morphology in the fetal dentate gyrus of E18 male, but not female, fetuses. In the placenta and fetal liver of males, inflammatory gene expression was affected by both DEP and maternal choline supplementation. However, maternal choline supplementation alone upregulated inflammatory gene expression in females, which may indicate an alteration in maturation rate. These data further contribute to the growing literature indicating region- and tissue-specificity in the developmental immune system in the context of maternal exposures.

## Introduction

Pollution exposure is a public health hazard, particularly during development. Air pollution from traffic and proximity to highways leads to developmental delays in brain connectivity and working memory capability (Pujol et al., 2016; Sunyer et al., 2015). Children who live in areas with high air pollution perform poorer on motor, IQ, and learning and memory tests than children in non-polluted areas (Wang et al., 2009; Calderón-Garcidueñas et al., 2011). In addition, prenatal exposure to polycyclic aromatic hydrocarbons, a common air pollutant, inversely correlates with widespread white matter alterations and behavioral problems such as ADHD symptoms, externalizing problems, and slower processing speed (Peterson et al., 2015).

It is possible to model human exposure to diesel particulate matter (a common pollutant in industrialized areas) within mice. One such model utilizes oropharyngeal aspiration to directly expose mice to precisely controlled levels of diesel exhaust particles (DEP). This model has been utilized to characterize the effects of prenatal diesel exposure in development. In this model, the adult mice whose dams were exposed to DEP show impaired hippocampal-dependent memory in male offspring only (Bolton et al., 2013). As well, deficits in anxiety-related behaviors and activity (Bolton et al., 2014; 2012) in DEP-exposed offspring compared to offspring that received maternal saline aspirations were observed. However, these effects are only “unmasked” with the addition of a second immune assault in adulthood.

Many of the effects of early-life pollution on behavior have been traced to neuroinflammation. Pollutant exposure *in vitro* leads to microglial stimulation of pro-inflammatory cytokines – specifically, interleukin-1β (IL-1β) and tumor necrosis factor (TNF, Sama et al., 2007). Though neuroinflammation has not been studied in human development as it pertains to air pollution, children do experience increases in asthma, an inflammatory disorder, due to increased air pollution (reviewed in Patel & Miller, 2009). Consistent with this increase in inflammation, findings in the aforementioned rodent model of early-life pollution have detailed male-specific inflammatory and behavioral alterations (Bolton et al., 2012; 2013; 2014; 2017), and there is a large body of work demonstrating the impact of maternal immune stressors on offspring development (reviewed in Knuesel et al., 2014; Bilbo & Schwarz, 2009). In particular, microglial “priming” is one mechanism by which early life inflammation impacts brain and behavior in adulthood: after an adult immune challenge, microglia overreact to the stimulus, leading to increased neuroinflammation and behavioral deficits (Bilbo, Biedenkapp et al., 2005; Bilbo, Levkoff et al., 2005; Bilbo et al., 2008). The lifelong deficits due to air pollution may be due to microglial priming.

Prenatal choline supplementation is neuroprotective throughout the lifespan. In a mouse model of autism, prenatal choline supplementation reduced anxiety-like behaviors of adult offspring (Langley et al., 2015). Prenatal choline supplementation also mitigates hallmarks of Alzheimer’s disease in aged offspring (Mellott et al., 2017) and the adult hippocampal memory deficits characteristic of Down syndrome (Velazquez et al., 2013; Moon et al., 2010). Prenatal choline supplementation also prevents age-related memory decline (Meck et al., 2008) and the decrease of hippocampal neurogenesis in advanced age (Glenn et al., 2008), indicating a lifelong effect and early-life “programming.” Prenatal choline supplementation is particularly effective in preventing adult neuronal and behavioral alterations due to seizure induction, such as cell loss (Wong-Goodrich et al., 2008), loss of neurogenesis (Wong-Goodrich et al., 2010), and impaired hippocampal learning and memory (Wong-Goodrich et al., 2010; Holmes et al., 2002).

Because choline is anti-inflammatory, prenatal choline supplementation may be able to blunt the immune dysregulation due to pollutant exposure early in life. To date, few studies have analyzed prenatal choline supplementation with a neuroimmune lens. In humans, circulating choline levels in pregnant women who experienced a recent infection were correlated with offspring protection in neurodevelopmental tests such as self-regulation and cerebral inhibition (Freedman et al., 2019), indicating that in humans, dietary choline prevents the behavioral effects of maternal immune activation. A study in a rodent model of maternal immune activation showed a protection in IL-6 expression in the fetal brain after a maternal immune assault (Wu et al., 2015). This landmark study provided evidence that the fetal immune system is impacted by maternal immune activation and maternal diet – neither of which were experienced directly by the fetus. Critically, dietary prenatal choline also normalized increases in adult autism-like behaviors due to maternal immune activation. Because of this work, we hypothesized that prenatal choline supplementation would similarly blunt the immune dysregulation in fetal brains, placentas, and fetal livers caused by prenatal diesel air pollution exposure in mice.

## Methods

### 1. Mice

Adult male and female C57BL/6 mice were obtained from Charles River Laboratories (Raleigh, NC, USA). Mice were time-mated using harem breeding of two females and a male. Upon confirmation of pregnancy with the visualization of vaginal plug (considered embryonic day 0, E0), females were pair-housed and given *ad libitum* access to water and the assigned diet. Specialized bedding (Alpha-Dri; Shepherd Specialty Papers, Milford, NJ, USA) was used to minimize risk of external contaminants. The colony room within the vivarium was on a reversed 12-hour dark-light cycle (lights off at 9am). These experiments were conducted with the approval of the Duke University Animal Care and Use Committee.

### 2. Prenatal manipulations

#### 2.1 Diesel Exhaust Particle (DEP) exposures

Briefly, diesel exhaust particles (DEP) were collected from a 4.8kW direct injection single-cylinder 320 mL displacement Yanmar L70V diesel generator at 3.5 rpm. Using an electrostatic precipitator, particles were collected from diesel exhaust. On the mornings of embryonic days 2, 5, 8, 12, and 16, dams were anesthetized with 2% isoflurane and exposed to diesel particles suspended in saline via oropharyngeal aspiration as previously described (Auten et al., 2012). Anesthetized mice were suspended by their frontal incisors on fishing line. Dams were exposed to either 50 µg of DEP dissolved in 50 µL of saline vehicle (phosphate-buffered saline [PBS] + 0.05% Tween 20), or vehicle alone. Using a 200 µL micropipette, DEP or saline solution was pipetted into the oropharynx by gently holding the tongue with forceps. Mice recovered from anesthesia under careful supervision. This method causes maternal lung inflammation similar to that seen after exposure to DEP via an inhalation chamber (Auten et al., 2012). This method was utilized over an inhalation chamber to administer the same amount of DEP in each installation and to each mouse; it is difficult to control diesel inhalation of mice in a chamber.

#### 2.2 Choline supplementation

At confirmation of pregnancy, dams were assigned to one of two diet conditions *ad libitum*: a synthetic control chow (1.1 g/kg choline chloride in formula AIN-76A with choline chloride substituted for choline bitartrate, DYET #110098, Dyets, Inc., Bethlehem, PA, USA), or the same diet with 4.95 g/kg choline chloride (DYET #110210; based on previous findings from Alldred et al., 2021; Maurer et al., 2021; Meck et al., 1988; Meck & Williams, 1997; Mohler et al., 2001; Powers et al., 2021; Velazquez et al., 2013; 2020).

### 3. Tissue collection

On embryonic day 18 (E18), dams were anesthetized with ketamine/xylazine (430 mg/kg ketamine; 65 mg/kg xylazine intraperitoneally, i.p.). Hysterectomy was performed to extract the fetuses, which were immediately placed on ice.

Fetuses were numbered and dissected in the order of their presence in the uterine horn from right to left. Whole heads were fixed in 4% paraformaldehyde. Whole placentas and one lobe of each fetal liver were flash frozen until analysis.

### 4. Fetal genotyping

To confirm the sex of each fetus, tails were genotyped for the *Sry* gene. This was done by DNA extraction with phenol, chloroform, and proteinase K. After DNA was obtained, the gene products of *Sry* were assessed by using polymerase chain reaction (PCR) and subsequent gel electrophoresis (Koopman et al., 1991). The primers used (Integrated DNA Technologies, Coralville, IA, USA) were as follows: forward (5’ – 3’): TGGGCTGGACTAGGGAGGTCC; reverse (3’ – 5’): TGCTGGGCCAACTTGTGCCT.

### 5. Fetal immunohistochemical analysis

#### 5.1 Iba1 immunohistochemistry

Fetal heads were cryoprotected in 30% sucrose after 48 hours in 4% paraformaldehyde, then gelatin-blocked. They were sliced using a cryostat at 14 µm and slices were thaw mounted directly onto slides in a series of 5. Briefly, slides that included the region of interest were washed with PBS before quenching with a solution of 50% methanol (VWR, Radnor, PA, USA) and 3% hydrogen peroxide (VWR) for 30 minutes. Slides were washed again in PBS and blocked in a solution of 5 mL 0.01M PBS, 150 µL normal goat serum (ThermoFisher Scientific, Waltham, MA, USA) and 50 µL Triton-X 100 (VWR) at 20°C for one hour and rinsed again in PBS. Slides were incubated at room temperature in a 1:500 concentration of rabbit anti-Iba1 (ionized calcium binding adaptor molecule 1) primary antibody (Fujifilm Wako Chemicals, Richmond, VA, USA). The next day, sections were washed in PBS and incubated for two hours in goat-anti-rabbit secondary antibody (Vector Laboratories, Burlingame, CA, USA) at a 1:200 concentration. Slides were washed and incubated in “Ready to use” Avidin/biotin complex (R.T.U. ABC, Vector). Sections were washed again and incubated in diaminobenzidine (DAB, SigmaFast 3,3’-Diaminobenzidine, Sigma-Adrich, St. Louis, MO, USA) until desired color was attained. Sections were washed again in PBS before they were mounted onto slides, dehydrated, and coverslipped.

#### 5.2 Unbiased stereology

Iba1+ cells were exhaustively counted in each target region using the optical fractionator method using StereoInvestigator software (Microbrightfield Inc., Williston, VT, USA). An optical dissector height of 7 μm, used a 50 μm x 50 μm counting frame, and counted cells using a 100x objective were used. Cells were counted only if the entire cell body was uniformly stained and had a minimum diameter of 13 μm (Kongsui et al., 2014). All brain regions analyzed were chosen because of previous research indicating an adult or fetal microglial morphological change in each area (Bolton et al., 2012; 2017). For each fetus, we analyzed the dentate gyrus (DG, Bolton et al., 2012; 2017), paraventricular nucleus of the hypothalamus (PVN, Bolton et al., 2012), basolateral amygdala (AMY, Bolton et al., 2012), and parietal cortex (PCX, Bolton et al., 2017). Five brain slices (including both hemispheres) per mouse were quantified for DG, PVN, and PCX analyses, and 3 per mouse for AMY analyses. Both sexes were quantified for all brain regions except for PCX, in which only males were quantified due to significant male microglial alterations in the other brain areas (Bolton et al., 2017). Iba1+ cells were classified into one of 3 morphological states based on cell shape, process thickness, and process number (Schwarz et al., 2012): round/amoeboid, stout processes, and thick long processes. The estimated number was divided by volume to calculate cell density. The density of “round” microglia, as well as density of all microglia, was analyzed.

### 6. Quantitative PCR

Quantitative PCR was utilized to assess the gene expression of *Tnf, Tlr2, Tlr4*, and *Itgam* in the placenta and fetal liver. Data values were excluded due to high Ct value (>32) or high technical replicate variability (>1.3 Cts). The Delta-Delta Ct method, in which the highest DCt of each group was subtracted from each DCt, was used to obtain a DDCt for each sample. Fold change was calculated as 2^(-DDCt)^.

### 7. Statistical analysis

The three variables of interest (sex, diesel particle exposure, and choline supplementation) were assessed using mixed-effects analysis, followed by Tukey’s post-hoc tests with Prism version 8.2.0 (GraphPad, San Diego, CA, USA). Because there were specific comparisons of interest, t-tests were also used to assess these effects. Statistical outliers identified with the ROUT method (Q = 10% for IBA1 results, Q = 1% for qPCR results) were excluded.

## Results

For the purposes of discussion, the following abbreviations will be used to refer to experimental groups: ConVeh (control diet + saline vehicle), ChoVeh (choline-supplemented diet + saline vehicle), ConDEP (control diet + diesel exhaust particles), and ChoDEP (choline-supplemented diet + diesel exhaust particles). Data are available here and on the Mendeley data repository (Maurer et al., 2022).

In the dentate gyrus of the hippocampus (DG), no significant differences between groups were observed using a 3-way ANOVA in either total Iba1+ cell density nor “round” microglial density (*p* > 0.05, Figure 1, raw data in Mendeley database). However, a t-test between ConVeh and ConDEP revealed a significant upregulation in total microglia (*p* =0.04) in the diesel-exposed males but not females. However, no effect of choline supplementation was observed.

**Figure 1.**
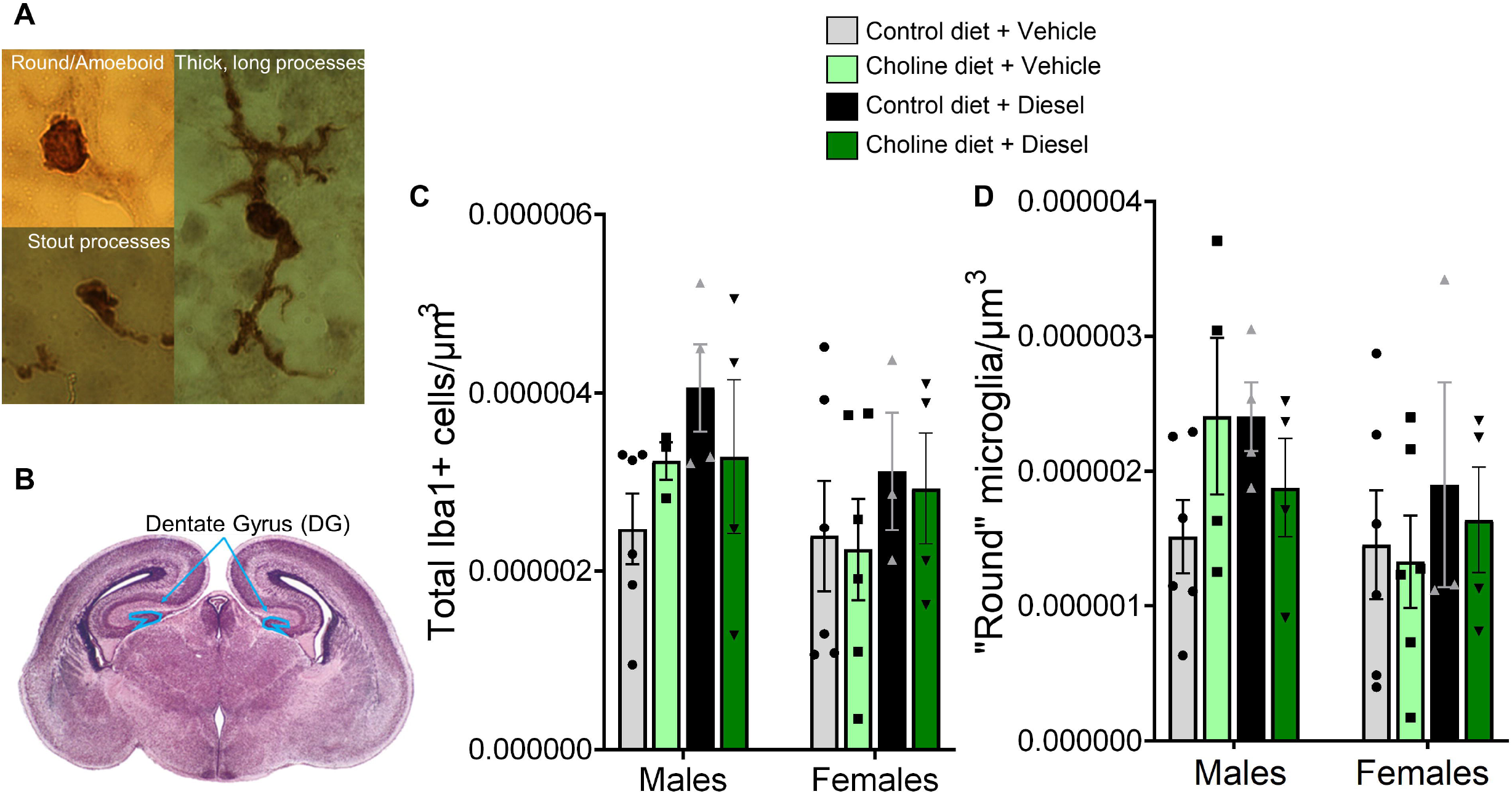
Maternal diesel exposure increases total microglial density in the male dentate gyrus. (A) Microglia were counted and classified into three morphological states. (B) Tracing of the E18 dentate gyrus, using an atlas image from Schambra, 2008. (C) No significant differences were found in the total density of microglia. However, a t-test revealed an increase in microglial density in ConDEP males only compared to ConVeh males. (D) No significant differences were found in the density of “round” microglia. N = 3-6 mice per group. Data are expressed as cell count/volume.

In the paraventricular nucleus of the hypothalamus (PVN), no differences due to maternal choline supplementation or diesel exhaust particles were observed in total microglial density (Figure 2B). Although no significant group differences in “round” microglial density were observed in a 3-way ANOVA, some marginal differences were observed. For example, across treatments males had more “round” microglia (*p* = 0.05, Figure 2C). As well, male ChoDEP brains had marginally more “round” microglia compared to male ConDEP brains (*p* = 0.09).

**Figure 2.**
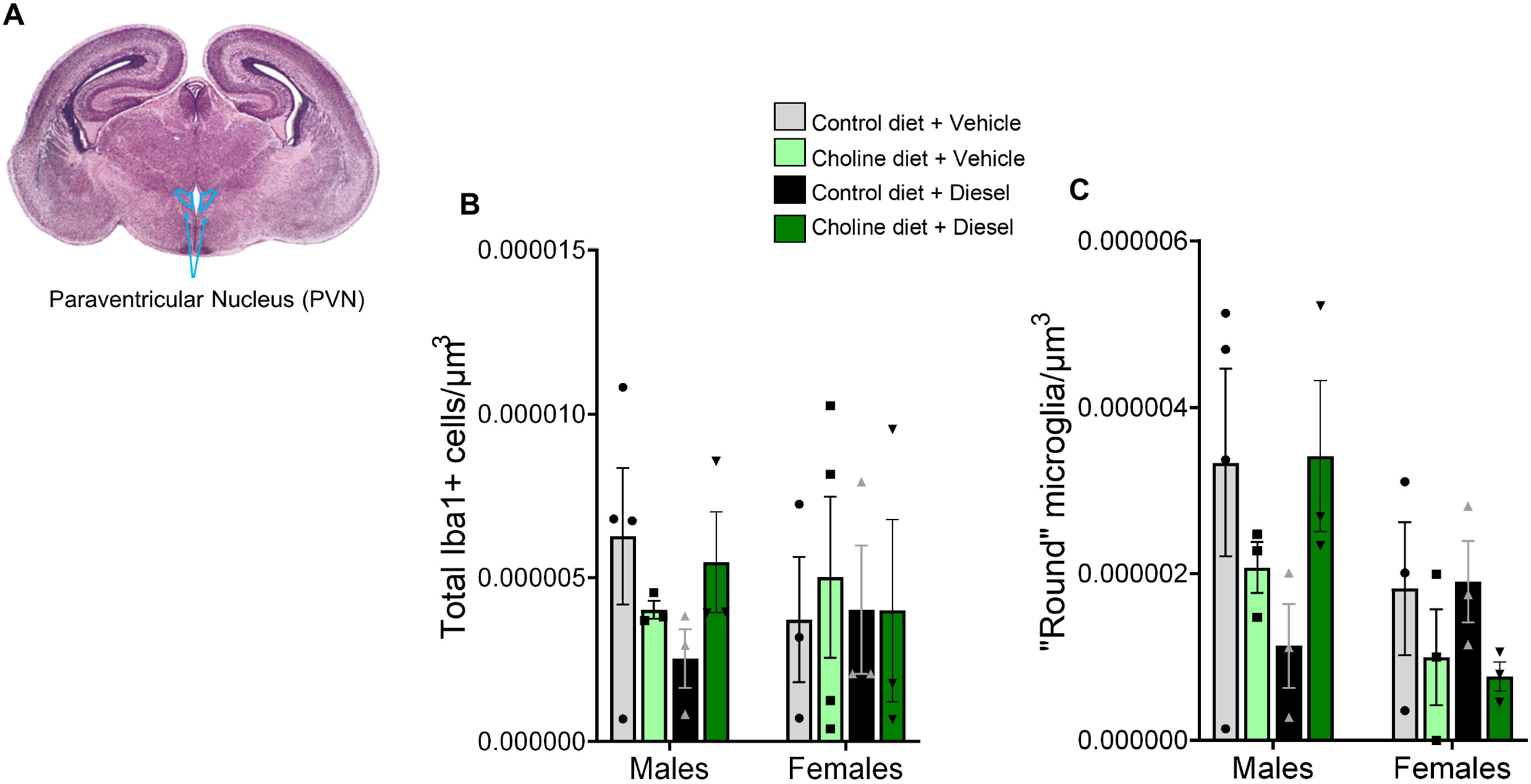
Maternal diesel exposure and diet had no significant effects on microglial density in the E18 PVN. (A) Tracing of the E18 PVN, using an atlas image from Schambra, 2008. (B) No significant differences were found in total microglial density in the PVN at E18. (D) No significant differences were found in the density of “round” microglia in the PVN. N = 3-4 mice per group. Data are expressed as cell count/volume.

In the amygdala (AMY), a significant 3-way interaction between sex, DEP, and choline supplementation was observed in total Iba1+ cell density (*p* = 0.03, Figure 3B) using a 3-way ANOVA. Although Tukey’s multiple comparisons test did not reveal any posthoc differences, t-tests in males only revealed that ChoDEP brains had significantly lower “round” microglial density compared to ChoVeh (*p* = 0.02) and ConDEP (*p* = 0.001). No differences in “round” microglia were observed in a 3-way ANOVA (*p* > 0.05).

**Figure 3.**
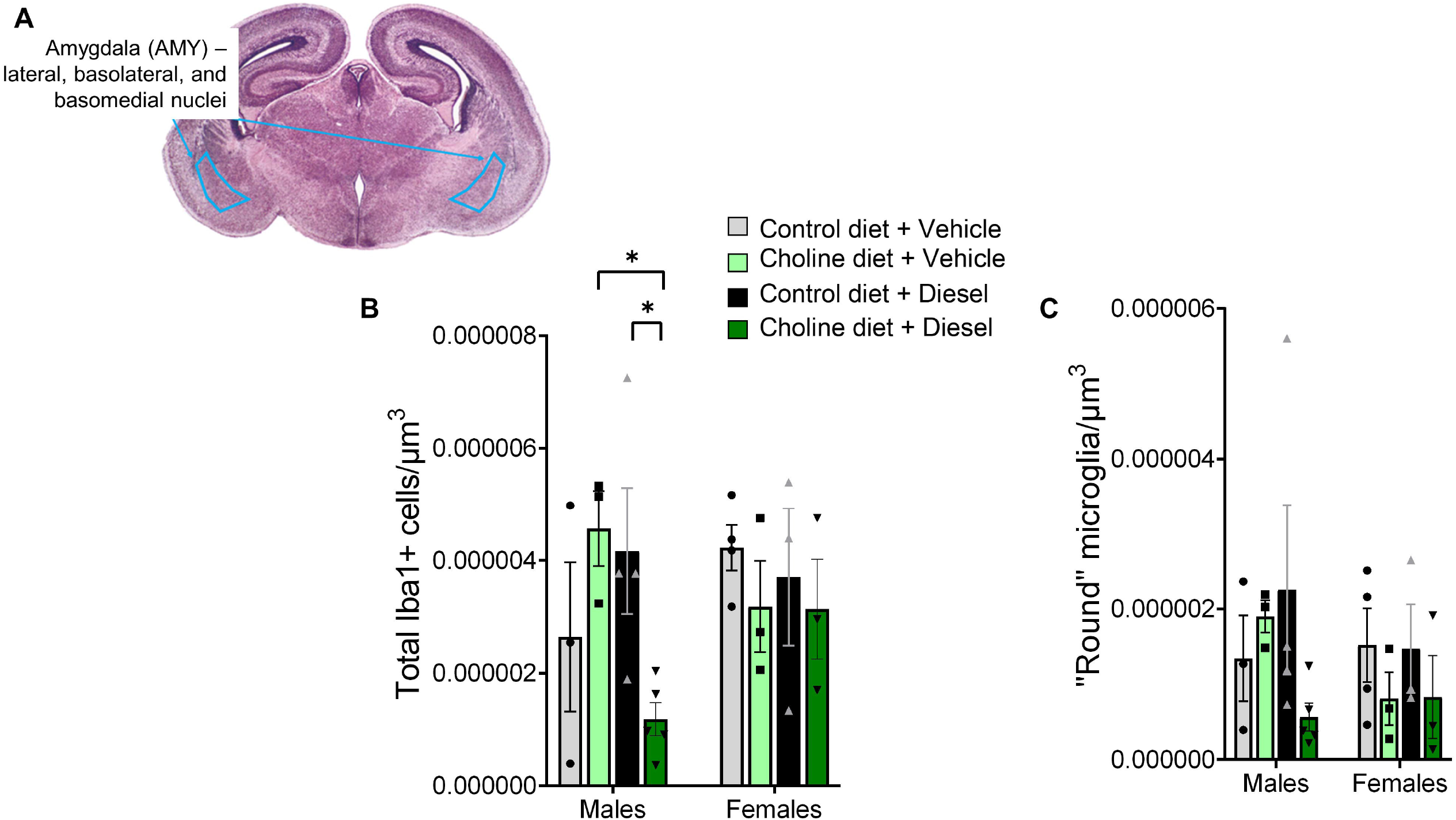
Maternal choline supplementation and diesel exposure significantly decreased the density of microglia in the AMY. (A) Tracing of the E18 AMY, using an atlas image from Schambra, 2008. (B) In the fetal male AMY, ChoDEP brains showed decreased “round” microglial density compared to ChoVeh and ConDEP males. (D) No significant differences were found in the density of “round” microglia in the AMY. N = 3-5 mice per group. \**p* < 0.05, t-test, males only. Data are expressed as cell count/volume.

In the parietal cortex (PCX, Figure 4), only male brains were quantified. No group differences were observed in microglial density or morphology in 2-way ANOVAs (*p* > 0.05).

**Figure 4.**
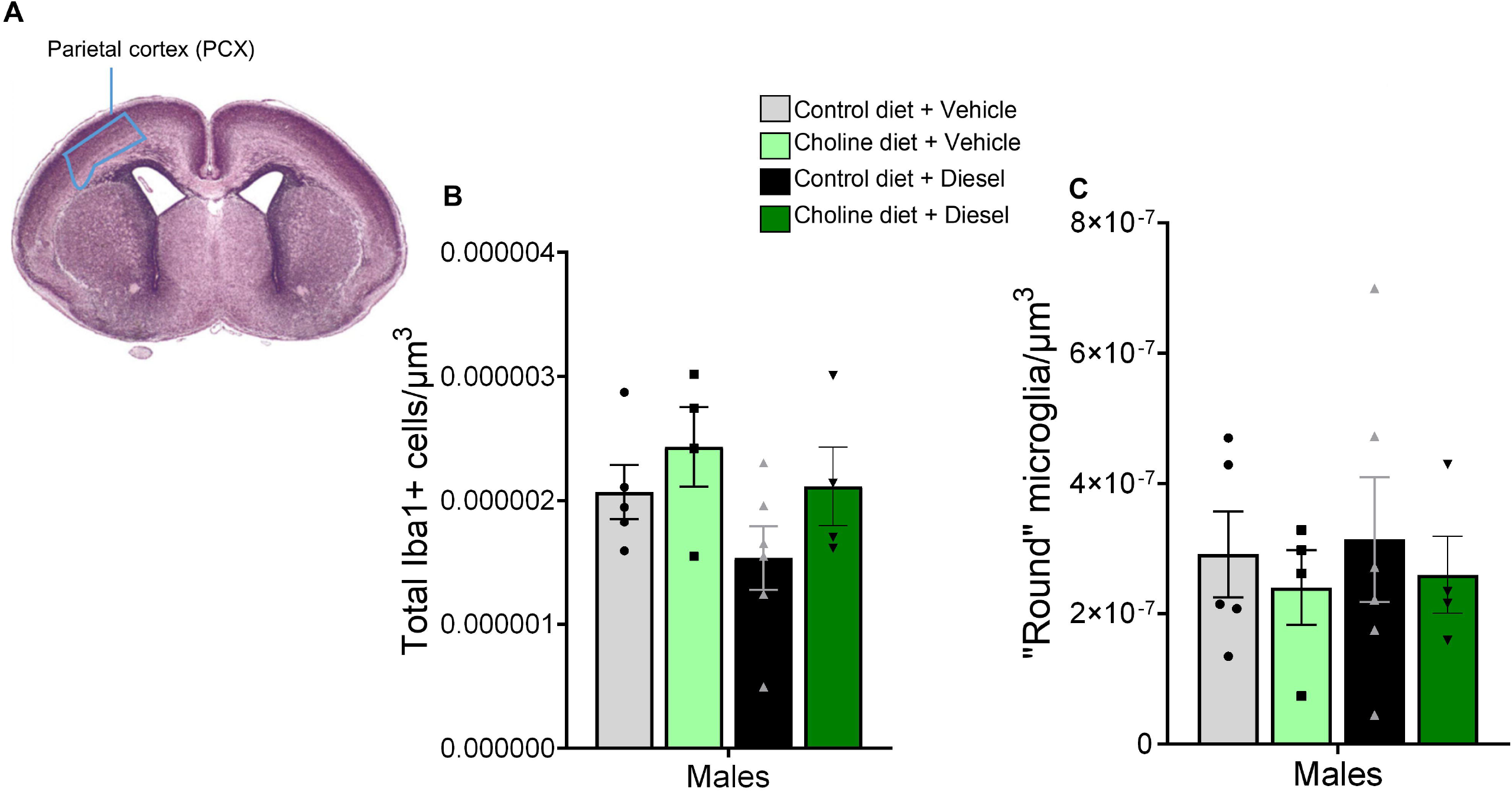
No effects of maternal choline supplementation or diesel exposure were observed in the E18 PCX. (A) Tracing of the E18 PCX, using an atlas image from Schambra, 2008. (B) No significant differences were observed in microglial density in the PCX. (D) No significant differences were found in the density of “round” microglia in the PCX. N = 4-6 mice per group. Data are expressed as cell count/volume.

Figure 5 reports the relative immune gene expression in the placenta. Regarding *Tnf* expression, a main effect of sex was observed in a 3-way ANOVA (Figure 5A, *p* = 0.03), with the placentas of female fetuses expressing more *Tnf* than those of male fetuses. A sex x choline interaction (*p* = 0.03) and a DEP x choline interaction (*p* = 0.01) were also observed. Tukey’s posthoc tests indicated that female ChoVeh placental *Tnf* expression was significantly higher than the expression of *Tnf* in male ChoVeh and ChoDEP placentas (*p* = 0.02, 0.03, respectively). In contrast, placental *Tlr2* expression was not significantly different between groups (Figure 5B).

**Figure 5.**
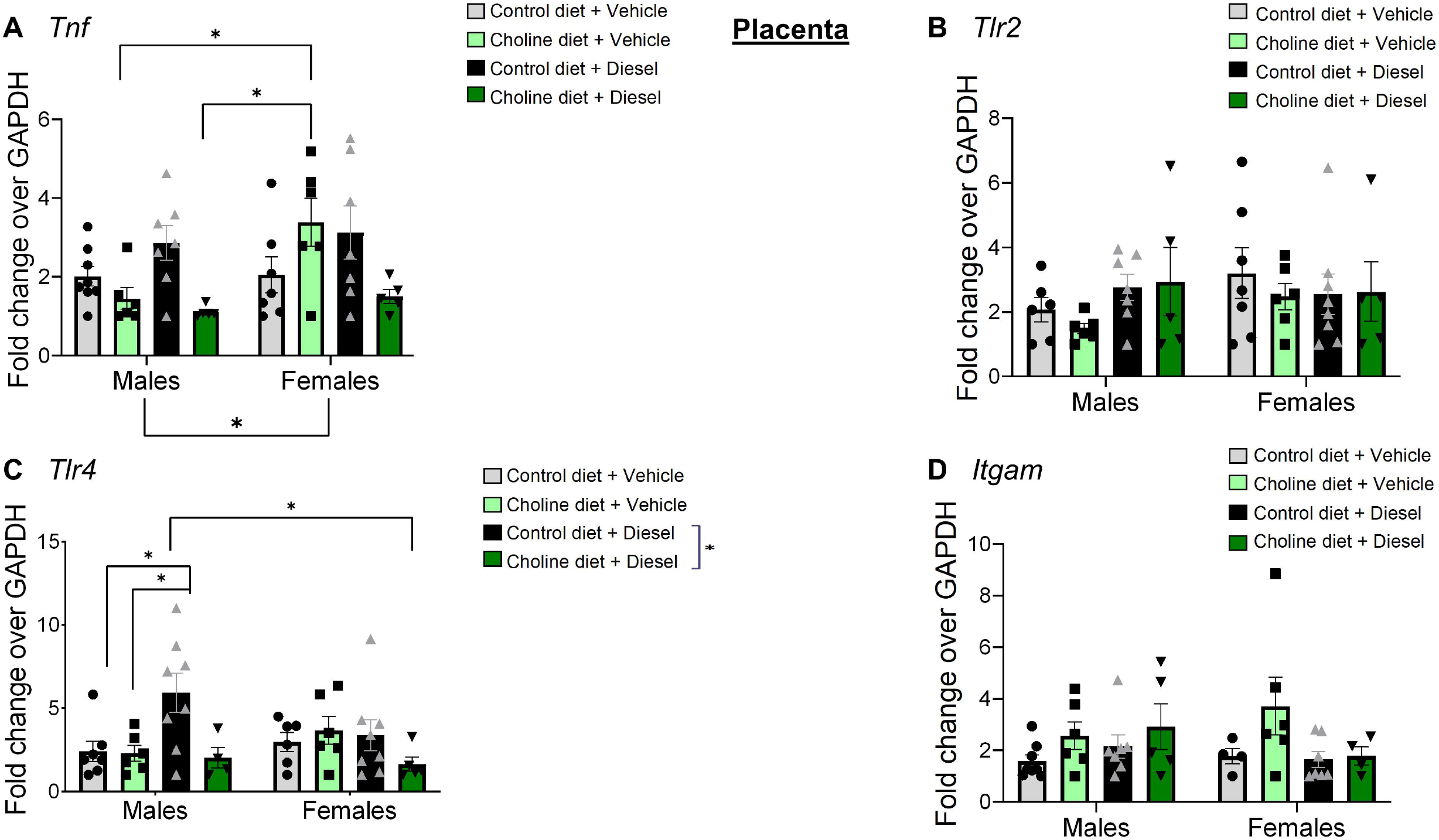
Inflammation-related gene expression in the placenta. (A) Females exhibit increased placental *Tnf*. (B) No significant differences due to diet or maternal diesel exposure in *Tlr2* expression were observed in the E18 placenta. (C) Maternal diesel exposure led to an upregulation of *Tlr4* in the male placenta. As well, in control diet groups only, DEP led to increased *Tlr4* expression in the placenta. Regardless of sex, ConDEP placental *Tlr4* expression was higher than in ChoDEP placentas. (D) Though a significant interaction was observed between sex and DEP, no posthoc differences were observed in placental *Itgam* expression. N = 4-8 placentas per group. \**p* < 0.05, Tukey’s multiple comparisons test. Data are expressed as fold change over *Gapdh*.

Placental *Tlr4* was also significantly different between groups. While no main effects were observed, two different interactions were significant: sex x DEP (*p* = 0.03) and DEP x choline (*p* = 0.03). Posthoc analyses revealed that diesel particles increased placental *Tlr4* expression in male placentas only. As well, control diet led to higher placental *Tlr4* in DEP-exposed placentas only. In the control diet groups only, vehicle treatment led to lower placental *Tlr4* expression than DEP treatment. Overall, regardless of sex, ConDEP placental *Tlr4* expression was higher than in ChoDEP placentas.

Placental *Itgam* expression was significantly altered via an interaction between sex and DEP, likely due to low expression in DEP-exposed females (*p* = 0.03, Figure 5D, raw data in Mendeley database). No posthoc differences were observed. No main effects of sex, choline or DEP were observed.

Figure 6 displays the relative immune gene expression in fetal livers. A DEP x choline supplementation interaction in liver *Tnf* was observed in a 3-way ANOVA (*p* = 0.02, Figure 6A). Within vehicle-exposed fetuses, choline supplementation led to an increase in fetal liver *Tnf* expression. No main effects were observed.

**Figure 6.**
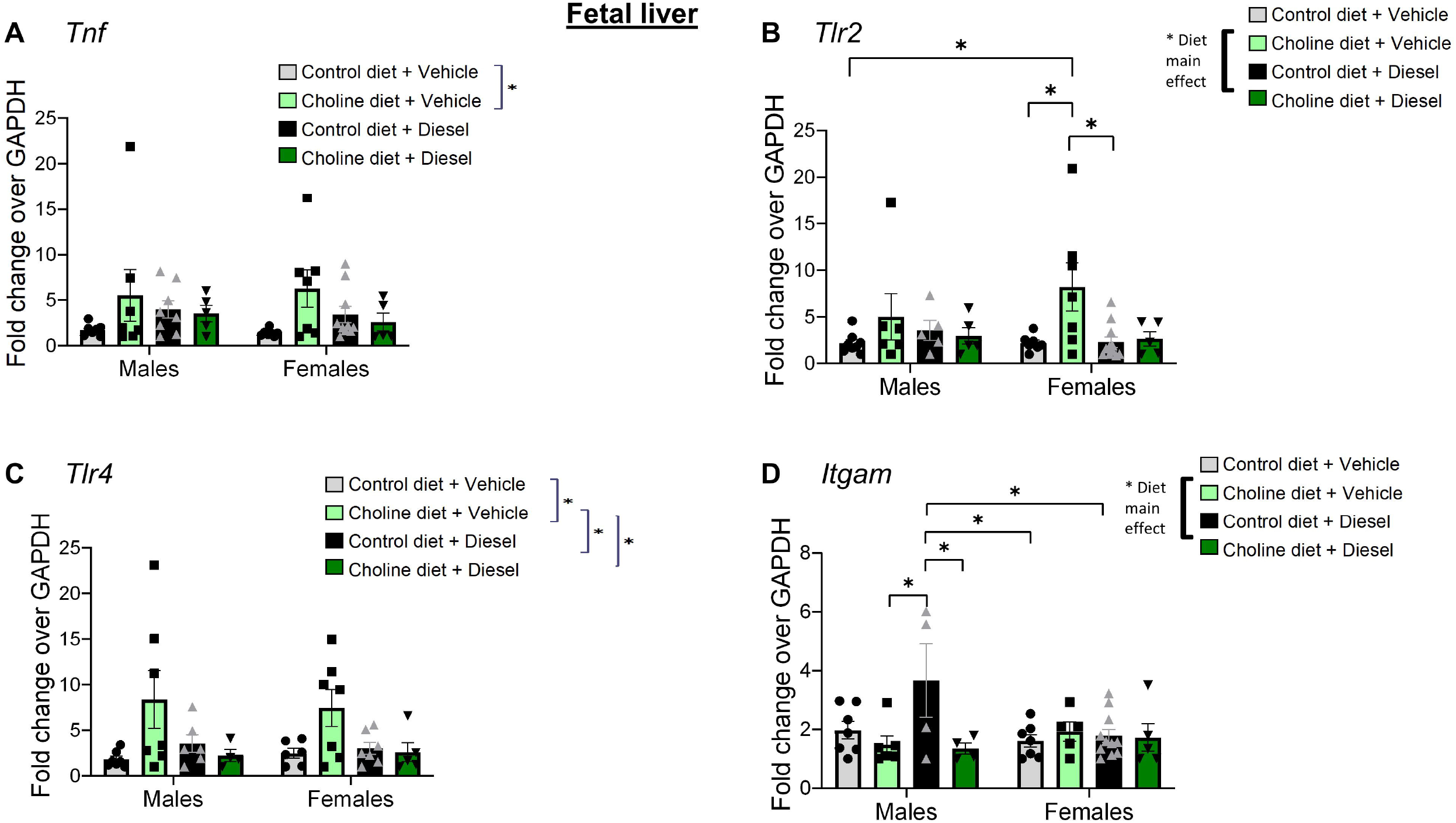
Inflammation-related gene expression in the fetal liver. (A) In vehicle-treated fetuses only, choline supplementation led to an increase in liver *Tnf* expression. (B) Maternal choline supplementation led to a significant increase in *Tlr2* expression in fetal female livers. (C) In the vehicle-treated groups, maternal choline supplementation led to a significant increase in fetal liver *Tlr4* expression. (D) DEP exposure led to an increase in *Itgam* expression in male fetal livers. N = 4-11 mice per group. \**p* < 0.05, Tukey’s multiple comparisons test. Data are expressed as fold change over *Gapdh*.

A significant main effect of choline supplementation was also observed in fetal liver *Tlr2* expression (*p* = 0.03, Figure 6B). As well, the interaction between DEP and choline was significant (*p* = 0.03). Posthoc analyses revealed that female ChoVeh fetal livers had higher *Tlr2* expression than male ConVeh, female ConVeh, and female ConDEP livers (*p* = 0.03, 0.046, 0.02, respectively).

While choline supplementation only marginally altered liver *Tlr4* expression (*p* = 0.08, Figure 6C), a DEP x choline supplementation interaction was significant (*p* = 0.03). Posthoc analyses revealed that ChoVeh fetal livers had higher *Tlr4* expression than all other groups.

Choline supplementation significantly decreased *Itgam* expression in the fetal liver (*p* = 0.04, Figure 6D). A significant interaction of sex x choline supplementation was also observed (*p* = 0.01). Posthoc analyses revealed that male ConDEP *Itgam* expression was significantly increased compared to male ChoVeh, male ChoDEP, female ConVeh, and female ConDep (*p* = 0.03, 0.03, 0.03, 0.04, respectively).

## Discussion

This work shows that, in the male fetal dentate gyrus, microglial activation was increased after chronic diesel exposure. Differences of this kind were only found in male fetuses, and not in other brain regions. Females were not affected by either diesel or choline in any brain region analyzed.

Maternal immune activation only leads to fetal immune activation in male mice; microglia of female mice do not appear to be altered by diesel or choline supplementation. Other research supports the view that male fetuses and not female fetuses are susceptible to immune activation caused by maternal DEP (Bolton et al., 2013; 2012; Ehsanifar et al., 2019). This male susceptibility to developmental delays due to maternal exposure to air pollution has also been shown in humans (Sears et al., 2019). Further supporting the hypothesis that males are impacted worse than females in models of early immune activation, early-life diesel exposure has been linked to male-specific increases in autism diagnoses in humans (Raz et al., 2018). Male and female fetal brains exhibit different inflammatory reactions to diesel exposure; specifically, male brains produce more IL-10 protein, but female brains exhibit a decrease in IL-10 protein (Bolton et al., 2013). This point is important because female brains are not necessarily resilient to DEP and maternal choline supplementation – they may be reacting differently than male brains, and these changes may not be detected by assessing microglial morphology.

The difference in microglial activation due to DEP in the fetal hippocampus in the present work may explain hippocampal-dependent behavioral deficits in adulthood. In another model of prenatal DEP, pollution dose-dependently impairs spatial memory in a Morris water maze (MWM) in the absence of a second immune assault in adult male offspring (Ehsanifar et al., 2019). Further, others using MIA models have found effects in adult hippocampal-dependent memory (Schaafsma et al., 2017) specifically in male offspring (Chlodzinska et al., 2011). The differences seen in microglial activation in the fetal hippocampus could possibly have large effects on behavior.

Taken together, the effect of maternal DEP and prenatal dietary choline supplementation is complex and appears to be both sex- and region-specific. Both diesel exposure and choline are systemic treatments, and do not work only on one tissue or cell type. These variables may cause changes in multiple biological pathways in addition to these neuroimmune changes. As well, the impact of these neuroimmune alterations on future behavior is not yet fully understood (Garay et al., 2013)

We have found that, in the dentate gyrus of the fetal hippocampus, prenatal DEP increases microglial density. Prenatal choline supplementation partially mitigates this activation increase. Only male fetuses were affected by diesel exposure. Clearly, both maternal inflammatory assaults and maternal diet impact the male fetal brain. Though the mechanism of maternal-fetal immune communication is not fully understood, DEP and choline supplementation have region- and sex-specific effects on microglia in the embryonic brain.

In the placenta, the most prominent finding was a male-specific increase in *Tlr4* due to DEP. Choline supplementation fascinatingly blocked this increase. *Tlr4* is expressed in placental tissue and immune cells, and its activation is associated with decidual inflammation (reviewed in Firmal et al., 2020). The role of TLR4 in the placenta, and its contribution to pregnancy complications such as preterm birth, preeclampsia, and gestational diabetes, has been well-reviewed (Firmal et al., 2020). Critically, all of these complications are also implicated in neurodevelopmental disorders (Crump et al., 2021; Maher et al., 2020; Wang et al., 2017).

Although the current set of experiments did not observe diesel-related increases in other placental immune markers, this particular increase could be indicative of the start of a vicious immune upregulation in the placenta due to diesel exposure. However, maternal choline supplementation mitigated this increase, providing support for its protective role in the placenta.

In the fetal liver, there was a male-specific increase in *Itgam* that was blunted by maternal choline supplementation. *Itgam* is the gene that encodes the CD11b protein. Recent work claims that fetal liver CD11b+ macrophages are “the major pro-inflammatory cells in the developing fetus” (Lakhdari et al., 2019), implying that this DEP-induced increase may be leading to more widespread fetal inflammation.

In conclusion, the effects of maternal DEP and maternal choline supplementation are complex and tissue-/brain region-specific in their actions. However, the current work seems to support previous studies showing male-specific impairments due to maternal inflammation. As well, in some measures, maternal choline supplementation can also blunt this upregulation – lending more support for its role as a neuroprotective nutritional supplement during pregnancy.

## Notes

https://data.mendeley.com/datasets/dn2cnhcnnm

### Competing Interest Statement

The authors have declared no competing interest.

